# Predicting transcription factor binding in single cells through deep learning

**DOI:** 10.1101/2020.01.14.905232

**Authors:** Laiyi Fu, Lihua Zhang, Emmanuel Dollinger, Qinke Peng, Qing Nie, Xiaohui Xie

## Abstract

Characterizing genome-wide binding profiles of transcription factor (TF) is essential for understanding many biological processes. Although techniques have been developed to assess binding profiles within a population of cells, determining binding profiles at a single cell level remains elusive. Here we report scFAN (Single Cell Factor Analysis Network), a deep learning model that predicts genome-wide TF binding profiles in individual cells. scFAN is pre-trained on genome-wide bulk ATAC-seq, DNA sequence and ChIP-seq data, and utilizes single-cell ATAC-seq to predict TF binding in individual cells. We demonstrate the efficacy of scFAN by studying sequence motifs enriched within predicted binding peaks and investigating the effectiveness of predicted TF peaks for discovering cell types. We develop a new metric “TF activity score” to characterize each cell, and show that the activity scores can reliably capture cell identities. The method allows us to discover and study cellular identities and heterogeneity based on chromatin accessibility profiles.

## Introduction

Transcription factors (TFs) bind to accessible or “open” promoter and enhancer regions and play a pivotal role in regulating gene expression by aiding or inhibiting binding of RNA polymerase(*1–3*). Different binding events lead to heterogeneity of gene expression across a population of cells, which may result in distinct cellular identities. Therefore, characterizing TF profiles of transcription factor is critical for understanding gene regulatory mechanisms and differentiating heterogenous cells.

Chromatin accessibility assays such as DNase-seq(*4*), Formaldehyde-Assisted Isolation of Regulatory Elements (FAIRE-seq)(*5*) and Assay for Transposase-Accessible Chromatin sequencing (ATAC-seq)(*6*) provide a way to study TF binding activity across the whole genome(*7*). Of these methods, ATAC-seq is gaining popularity due to its comparative cost-efficiency and simplicity. ATAC-seq profiles are generally designed to identify open chromatin regions, which can be used to infer TF binding events where these regions overlap with protein binding sites. Indeed, methods such as FactorNet(*8*) and deepATAC(*9*) leverage a deep learning-based approach to identify open chromatin regions and infer TF binding locations using bulk chromatin accessibility data. However, both of these methods are only applicable to bulk data, and therefore do not take into account heterogeneity within cellular populations.

Recent advances of single cell epigenomic sequencing permit characterization of chromatin accessibility at a single cell level(*10*). For example, probing chromatin accessibility within single cells by scATAC-seq has become possible(*11–13*), enabling the identification of *cis*- and *trans*-regulators and the study of how these regulators coordinate in different cells to influence cell fate(*13–15*). Unfortunately, as with all single cell transcriptomics, scATAC-seq data are sparse and noisy due to not only technical constraints such as shallow sequencing(*11*) but also biological realities such as cellular heterogeneity(*16*).

In order to address these challenges, we present a deep learning-based framework called single cell Factor Analysis Network (scFAN). scFAN pipeline consists of a “pretrained model”, which is trained on bulk data, and is then used to predict TF binding at a cellular level using a combination of DNA sequence, aggregated similar scATAC-seq data and mapability data(*17*). This approach alleviates the intrinsic sparsity and noise constraints of scATAC-seq. scFAN provides an effective tool to predict different TF profiles across individual cells and can be used for analyzing single cell epigenomics and predicting cell types.

## Results

### scFAN overview

We start with a brief overview of scFAN (Fig. 1a). scFAN is a deep learning model, receiving ATAC-seq, DNA sequence and DNA mapability data at a given genomic region as input, and predicting the probability of a TF binding to this region. scFAN is trained using publicly available “bulk” datasets, which contain genome-wide ATAC-seq and ChIP-seq profiles collected from multiple cell types measured at a population level. The data inputs (i.e. input feature vectors) are 200 bp bins composed of bulk ATAC-seq data, DNA sequence and mapability data for that bin. The feature vectors are fed into a 3-layer convolutional neural network (CNN), which is then linked to 2 fully connected layers and a final sigmoid layer to make predictions. The ground-truth outputs are binary labels indicating TF binding, annotated based on ChIP-seq peaks.

**Figure 1:**
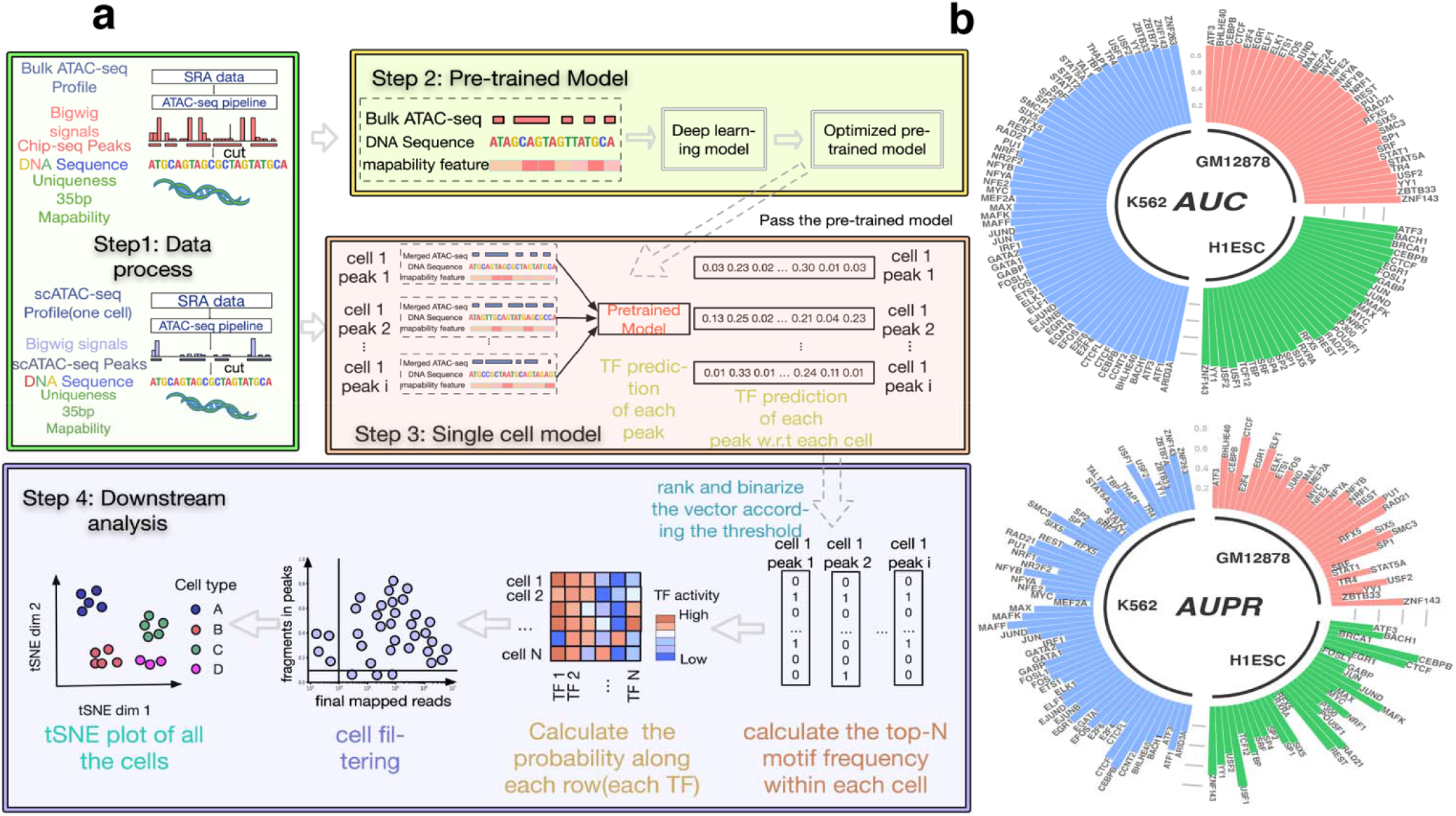
scFAN shows strong classification performance. **a** scFAN pipeline. Bulk ATAC-seq, mapability data and regions of DNA identified by ChIP-seq data is passed to the deep-learning “pre-trained model”. The trained model is then used to predict TF binding profiles based on regions of DNA called by scATAC-seq, mapability data, and a combination of scATAC-seq and bulk ATAC-seq. TF “activity scores” are calculated from the predictions by summing the number of times the top two most frequent TFs appear per cell. scFAN clusters cells from these activity scores. **b** Circular barplots showing AUC and auPR values of all the TFs from the pre-trained model, from 3 different cell lines.

Once the model is fully trained, scFAN predicts TF bindings in each individual cell based on scATAC-seq profiles measured in that cell. Due to the intrinsic sparsity of current scATAC-seq technology, we first impute the signal of scATAC-seq at individual bases of each cell by aggregating scATAC-seq data from similar cells. For each single cell, we calculated similarity scores between it and other cells and aggregated chromatin accessibility signals of its similar neighbors to boost chromatin accessibility coverage and used the aggregated data as inputs to our model. This approach allows us to increase the chromatin accessibility coverage while retaining cellular specificity.

The input vectors in the prediction step are the aforementioned aggregated scATAC-seq data, DNA regions called by scATAC-seq, and mapability data.

### Validation of scFAN accuracy on bulk data

We trained scFAN on three bulk ATAC-seq datasets, GM12878, H1-ESC and K562, in which ChIP-seq data for a number of TFs are also available from the ENCODE consortium (with 33, 31 and 60 TFs in each dataset respectively), and generated three pre-trained scFAN models – one for each dataset. We then validated the accuracy of the trained models on test datasets (hold-out chromosome regions not used during training). Similar to the TF binding annotations in the training data, the ground-truth labels of the TF binding in the testing data are also based on ChIP-seq peaks. The prediction accuracy was measured by the Area Under the ROC Curve (AUC), the Area Under the Precision-Recall curve (AUPR) and the Recall value corresponding to each TF (Fig 1b, Supplementary Fig S3, S4). Our trained model captured most of the TF binding information correctly: all the TF prediction AUC values are over 0.80 and nearly half of the TF AUPR values are over 0.8 (Supplementary Table S1). What is more, we and others have reported that convolutional neural networks could capture TF binding motifs information(*8, 18*). We used the same heuristic from FactorNet and visualized TF kernels of SPI1, CREB1, JUND and MAFK from the trained model based on cell line GM12878. These kernels were first converted to position weight matrices and then aligned with motifs from JASPAR(*19*) using TOMTOM(*20*). All these kernels successfully matched the TFs that were identified by known database like JASPER with matched E-values all less than 10^-3^, e.g. 9.02*10^-4^ for TF SPI1 (Fig. 2b).

**Figure 2:**
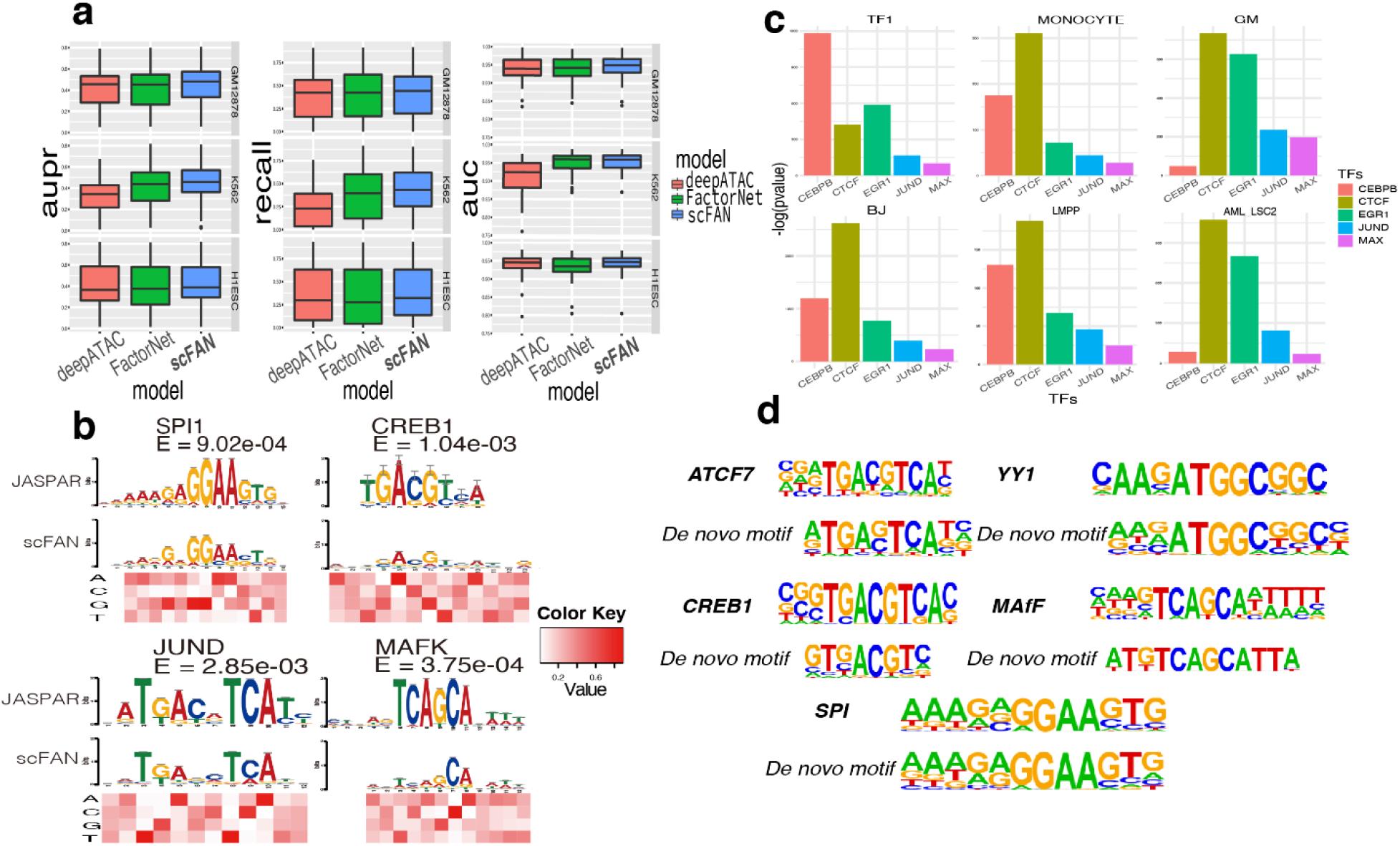
Validation on TF predictions. scFAN can predict both bulk and single cell TF binding. **a** Box plot of the performance of the pre-trained model and two other models predicting bulk cell TF binding on the same dataset. **b** Four convolutional kernels that matched with four known motifs derived from JASPAR database. The heatmap denotes the value of each nucleotide corresponding to above position. **c** Enrichment analysis of the 5 predicted mostly active TFs from 6 randomly chose cells. scFAN predicts the most likely TF per bin, adds up the number of times each TF is the highest predicted TF. Homer takes all the candidate peaks need to be predicted and generates the enrichment analysis. All these TFs were significantly enriched in all these peaks. **d** Separately selected regions predicted by scFAN whose most active TF are ATCF7/YY1/CREB1/MAfF/SPI were performed enrichment analysis, de novo matched motifs were compared to the known motifs from Homer.

We then compared two other state-of-the-art bulk TF binding profile prediction methods to scFAN, FactorNet(*8*) and deepATAC(*9*). Similar to FactorNet and deepATAC, scFAN uses convolutional neural nets as its basic building structure but simplifies the model structure to include fewer convolution layers with fewer parameters. A key difference between scFAN and the previous two models is scFAN’s ATAC-seq signal is continuous, as opposed to the binarized signal used by FactorNet and deepATAC. Binarizing the data may result in loss or change of ATAC-seq signal depending upon how the data are binarized. To make the comparison fair, all three models were trained and tested on the same datasets. Encouragingly, scFAN more accurately predicted bulk TF binding than either FactorNet or deepATAC, based on mean values of AUC, auPR and Recall in three cell lines (Fig. 2a). Per the GM12878, K562 and H1ESC cell lines, 85% (61%), 90% (55%) and 81% (71%) of TF predictions have better recall values compared with deepATAC (FactorNet). The improvements are statistically significant for two comparisons (two-tailed t-test, *p* < 0.05).

### Single cell TF predictions are consistent with enrichment analysis

Next, we evaluated scFAN’s predictive performance at a single cell level. We ran scFAN TF binding prediction on a scATAC-seq dataset consisting of 2210 cells with a mixture of cell types ranging from the chronic myelogenous leukemia cell line K562 of drug treated and untreated to lymphoblastoid cell lines (LCLs; GM12878) (including replicates), human embryonic stem cells (H1ESC), fibroblasts (BJ), erythroblasts (TF-1), promyeloblast (HL60), acute myeloid leukemia (AML) patients, lymphoid-primed multipotent progenitors (LMPPs), and monocytes cells from Corces *et al*(*11, 21*). For each individual cell, we ran TF binding prediction using each of the three pre-trained scFAN models and then concatenated these predictions (See Methods) to generate the binding profiles of 88 TFs in each of the 2210 cells.

Unlike the bulk data, TF information in ChIP-seq at a single-cell level is not possible at this stage due to technical constraints. Hence, we cannot evaluate the accuracy of our single-cell TF binding predictions by directly comparing to a ground-truth label as in the case of the bulk data. To assess the quality of our predictions, we instead use two indirect approaches.

First, we verified whether there are sequence motifs enriched within the predicted TF regions and whether these motifs match known binding profiles of the TFs. For this purpose, we used the software Homer to discover and evaluate the enrichment of motifs with scFAN predicted peaks(*22*). The result showed that TFs of five active TFs predicted by scFAN in six cell types were all significantly enriched in Homer (*p* < 10^-10^, Fig. 2c). Intriguingly, TFs critical to monocyte differentiation such as SPI1 (a.k.a. PU.1), EGR, CREB and YY1 were highly enriched in monocyte cells (p-value < 1e-5)(*23, 24*). To further explore whether each of these TF binding prediction matches the known motifs, we implemented TF prediction in all the candidate peaks in the monocyte cell using scFAN, then selected those peaks that were predicted to bind with each one of these TFs, and finally performed *de novo* enrichment analysis using Homer. For each TF result, we used the one of the most enriched *de novo* assembled motifs to match with corresponding TF motifs, and we found that the *de novo* assembled motifs from scFAN closely matched known motifs from Homer (Fig. 2d).

### Using single cell TF prediction to cluster cell types

Next, we studied whether the predicted TF binding profiles can be used to differentiate cell types. We reasoned that if the TF binding predictions are accurate, it should be able to use them to cluster cells into different groups that share similar cell identifies. Fortunately, the cell types of individual cells in the scATAC-seq datasets are known. We can therefore assess the quality of the cell clusters derived from TF binding profiles by comparing them to true cell type labels.

To explore scFAN’s ability to cluster cell types based on the TF binding predictions, we developed a new metric called “TF activity scores” to characterize the state of single cells. The TF’s activity score of a cell summarizes the intensity of its predicted occurrences across the genome in the cell – the higher the score, the more active the TF is (See Methods). Overall the state of each cell is characterized by a TF activity vector of dimension 88, one component for each TF (all three pre-trained models were used to generate TF activity scores, merged together with 88 components. See Methods). The 2210 cells are clustered using K-means clustering based on Euclidean distances between the TF activity vectors, shown in tSNE plots (Fig. 4a).

To validate the clustering result, we evaluated the clustering performance of scFAN by comparing the predicted clusters to ground-truth cell type labels. The performance of scFAN was benchmarked against several other popular methods that cluster cells on chromatin accessibility data – scABC(*25*), cisTopic(*26*), SCALE(*27*), Cicero(*28*), Brockman(*29*) and ChromVAR(*14*). We performed the same filtering procedure for all the cells and used the same parameter settings (Supplementary Fig S2). We used three common metrics to quantitatively measure the clustering performance of scFAN and other compared methods: Adjusted Rand Index (ARI)(*30*), Normalized Mutual Information (NMI)(*31*) and V-measure score(*32*). Our model had the highest metric scores among all these methods with ARI, NMI, and V-measure score equaling 0.470, 0.674, and 0.674 respectively (Fig. 3b). These results indicate that clustering cell types based on TF activity scores is consistently better than previous methods based on peak-cell matrix or chromatin accessibilities.

**Figure 3:**
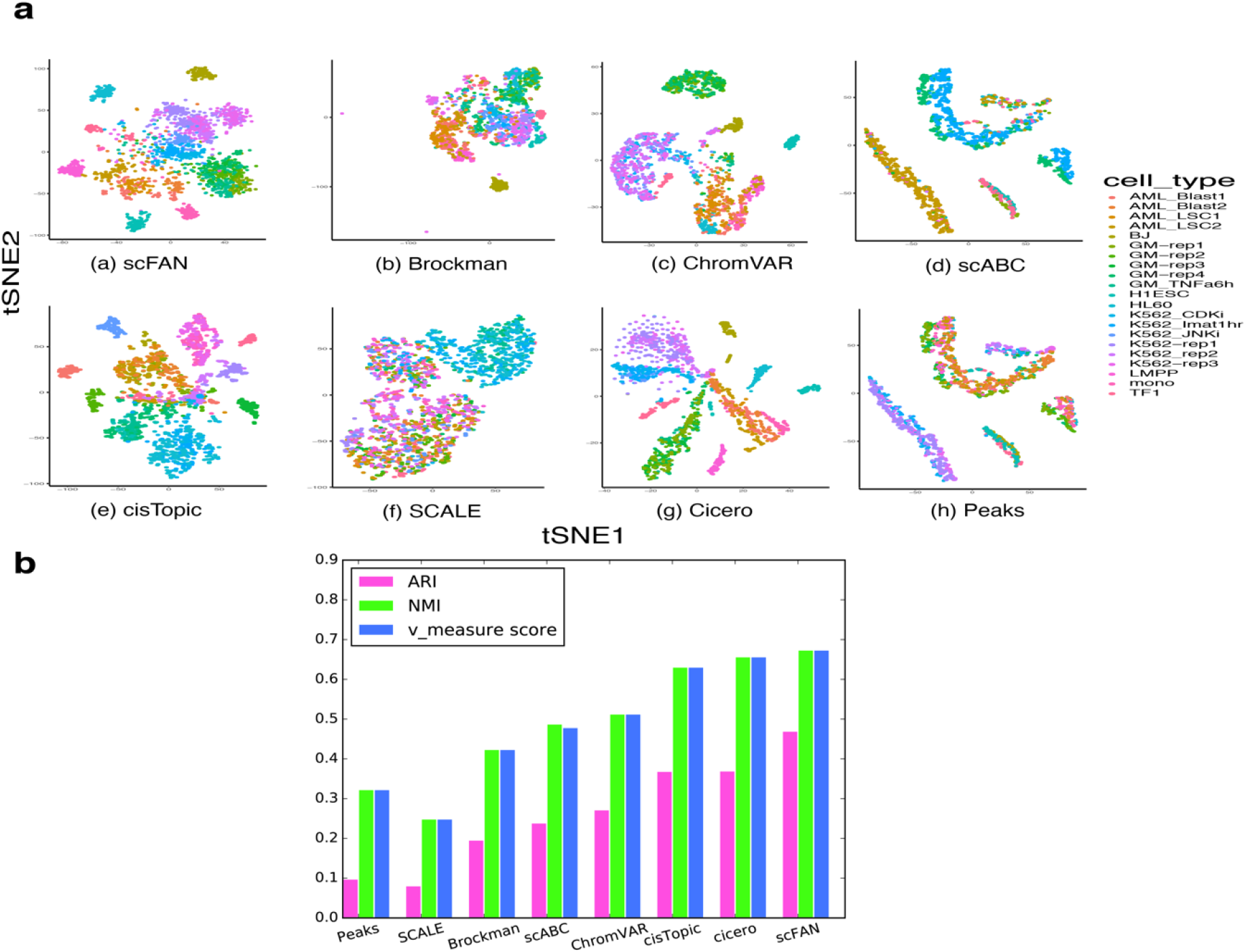
Comparison of scFAN to 7 other count matrix based methods or open chromatin accessibility-based methods. **a** tSNEs of all seven different open chromatin-based or count matrix based clustering methods. **b** Comparison of 7 different clustering metrics of each method. ARI, NMI, and v-measure score were used to measure each method. The higher the score is, the better the clustering performance is.

Having demonstrated that TF activity scores are effective in differentiating cell types, we explored the contribution of individual TFs in defining cell identifies. For this purpose, we plotted the activity scores of three TFs (EGR1, CEBPB and SPI1) across the 2210 cells on top of the cluster tSNE plots (Fig 4a). A couple of observations are notable from these plots. First, individual TF shows considerable amount of variations in its activity scores across different cell types. For instance, EGR1 is highly active in LMPP cells with a mean activity score of 1.879, higher than any other cell type, suggesting its prominent role in the transcriptional regulation of LMPP cells. And so is CEBPB in BJ cells (Fibroblast), which has the highest mean activity score of 3.136 over other cells. Second, there is also large heterogeneity among different TFs in their involvement in different cell types. SP1 is more active than EGR1 in monocyte cells, with EGR1 mean activity score value higher in AML cells than monocyte cells and the converse for SPI1. These observations seemed to be consistent with previously published studies which not only indicate that EGR1 is highly enriched in LMPP cells(*33*), but also show that CEBPB is more involved in Fibroblast cell development(*34*) and so does SPI1 in LMPP cells(*35*).

**Figure 4:**
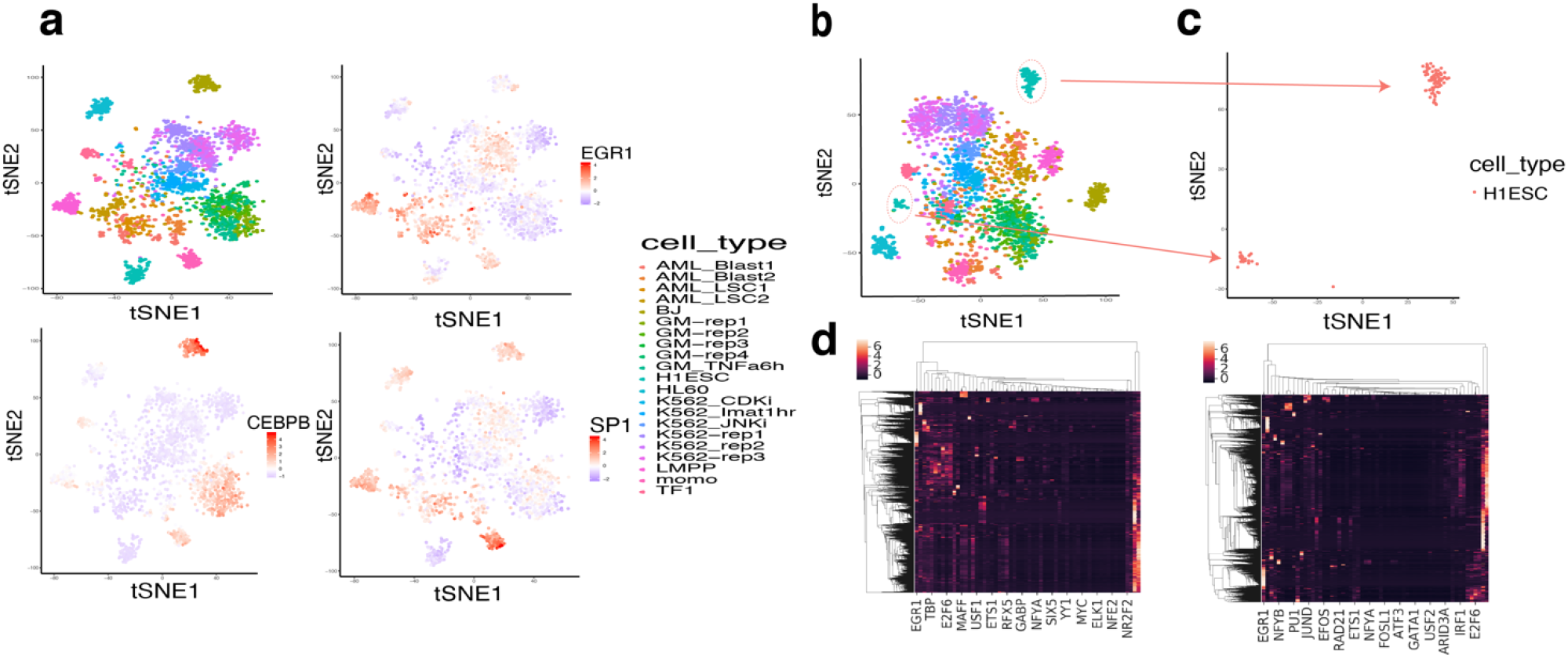
H1 cells are well separated by TF activity scores. **a** TFs have varying activity scores across cell types, EGR1 is more involved in the LMPP cells, CEBPB is more involved in BJ(Fibroblast) cells, and SPI1 is more related to monocyte cells. **b & c** Separation of H1 embryonic stem cells (ESCs) colored red when using scATAC-seq as model input, the H1ESC cells clearly separated into two distinct groups (subcluster 1 and 2). **d** Heatmap plot of all the TFs and across the whole chromosome from one H1ESC cell, left is using aggregated scATAC-seq data as input, right is using the raw scATAC-seq data as input. The left heatmap contains more TF prediction information than the right plot.

Using scFAN and TF activity score-based clustering can potentially help alleviating single cell sparsity and improve clustering performance. When only raw scATAC-seq data without aggregation was used to predict TF and clustered cells, scFAN subclustered nominally genetically identical H1ESCs (Fig 4b,4c). However, when we adopted the aggregated scATAC-seq data as our input, scFAN managed to drag the two subclusters back together into one cluster. It showed that the aggregation of the scATAC-seq signals probably helped to recover chromatin accessibility signals of H1ESC cells, which made the model prediction be more accurate. The heatmap plots of TF prediction on one H1ESC cell across all the peaks using raw scATAC-seq data and the aggregated scATAC-seq data showed that the TF prediction results of scFAN contain higher probability on some TFs compared with the heatmap without scATAC-seq aggregation, which has brighter color in the figure (Fig 4d). Furthermore, we randomly selected the regions in chromosome 1 to visualize the chromatin accessibility signals (Supplementary Fig S5-S6). And we found some regions that are lack of signal became dense after borrowing information from neighboring cells, which might be helpful for the further TF prediction in those separated H1ESC cells. From the improvement on ARI and NMI metric values we could indirectly infer that scFAN potentially has the ability to help alleviate the data sparsity and find some missing signal in scATAC-seq data within some low coveraged single cells and thus provide a better performance on TF prediction across the genome.

### Performance and sensitivity

Since we’ve shown that scFAN can cluster cells accurately, we wanted to characterize how sensitive the clustering is to different parameters. We started by varying the number of top predicted TFs per cell the clustering takes into account. scFAN by default uses the top 2 predicted TFs per cell. We compared the original clustering result to clustering based on the activity scores of only the most active TF and also the top 5 most active TFs. There is a slight improvement in the NMI and v-measure when choosing the top 5 TFs, but top2 yields the highest ARI score, overall the clustering is robust to the chosen number of most active TFs (Fig. 5a).

**Figure 5:**
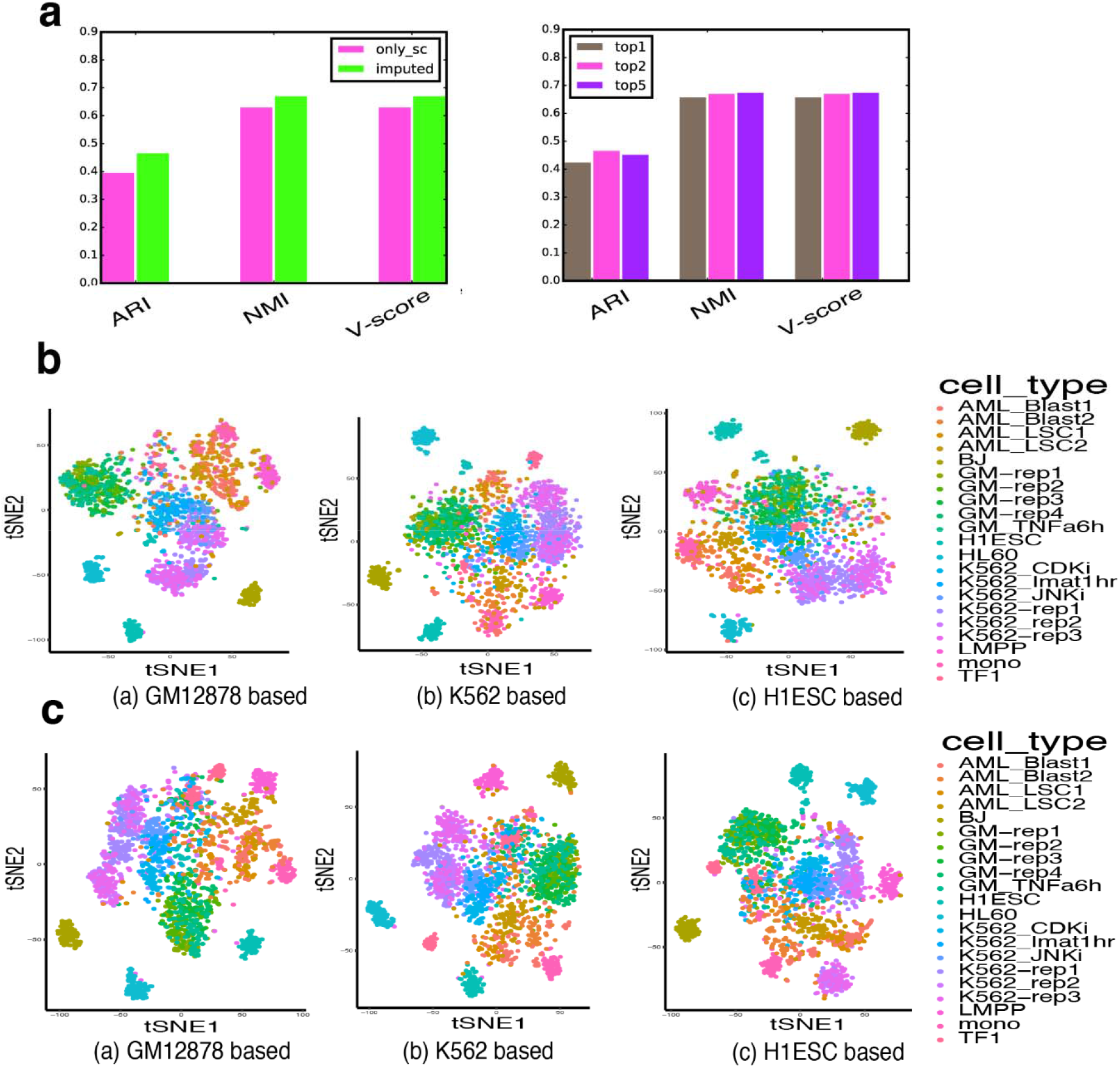
Clustering performance comparison when different thresholds and parameters are changed. **a** Sensitivity of clustering influenced by different ATAC-seq that is been used in the model and the Sensitivity of clustering to number of top TFs used. scFAN by default clusters cells on the combined ATAC-seq data and on the activity score of the top two most active TFs. **b** Clustering performance using 3 different pre-trained models adopting scATAC-seq data as input. **c** Clustering performance using 3 different pre-trained models adopting aggregated ATAC-seq data as input.

We also predicted clustering with just raw scATAC-seq data. As before, the pre-trained model was the same in both the default case but just used scATAC-seq data without imputing signal. We found that including similar single cell ATAC-seq data to alleviate the data sparsity drastically improves clustering performance over only using scATAC-seq data (Fig 5a, 5c), probably due to the aformentioned sparsity and noisiness of scATAC-seq.

## Discussion

Here we developed what is, to our knowledge, the first pipeline to predict TF binding at a cellular level. scFAN is a deep learning-based single cell analysis pipeline which mitigates the fundamental difficulties in analyzing scATAC-seq by leveraging bulk ATAC-seq data. We found that scFAN can predict TF binding motifs accurately. scFAN robustly identifies cellular identities, even in cells that are genetically similar. Detecting cellular identities at a chromatin accessibility level may enable more faithful and identification of distinct cell types.

Since the limited number of datasets used in this study lack full coverage of all TFs in humans, some TF activities may be missing. With availability of more TF related data covering more TFs across multiple cell types, merging such TF related single cell information into one dataset could lead to better prediction of TF binding and avoid calibration on prediction results, which is also a further expectation of implementing scFAN to more data. The pre-processing time, e.g. binning the peaks, was found to be variant based on the number of peaks: for instance, two minutes were needed to bin a cell with over 10,000 peaks. The pre-processing time makes the total time from preprocessing to final prediction highly variable for different cells, and parallelizing such pre-processing of each cell will speed up the prediction. The speed of training for the pre-trained model usually is less critical because the trained model can be reused on different datasets. Since scFAN is the first method of such kink, a lack of other single cell TF binding profile prediction methods or established benchmarks makes validation more challenging. While comparing predicted motifs to previously known ones or clustering cells based on TF activity can provide validation to some degree, experimental validations need to be carried out, for instance, using single cell ChIP-seq(*36*) for further confirmation on the predictions made by scFAN. Last but not the least, more and effective downstream analyses, such as psuedotime analysis, can also be further expanded and refined in scFAN.

Overall, scFAN is a highly promising tool for single cell analysis, not only for predicting TF binding and TF motifs but also for determining cellular identities. Being able to correlate open chromatin regions and binding activity of transcript factors in individual cells enables better understanding of cellular dynamics and regulations. This study shows that the deep learning technique can significantly improve our capability of utilizing single-cell data to discern cell fate decisions.

## Materials and Methods

### Data processing

#### DNA sequence

The sequence data was processed as in Quang *et al*(*8*). The genome was segmented into 200 bp bins, containing both the forward and reverse strands, with 50 bp intervals. Bins that overlapped with a known TF binding site were considered “bound” bins, whereas bins that overlapped with the blacklist region(*37*) were labeled “unbound” bins. Bins that overlapped with neither region were discarded. The bins were then expanded to 1000 bp, centered around the middle of each bin locus.

#### Chromatin accessibility

We processed the raw bulk ATAC-seq files by trimming with cutadapt(*38*) mapped to the human genome(hg19) using Bowtie2, and discarded the redundancy read pairs using Picard. We processed the scATAC-seq data with the ENCODE ATAC-seq pipeline protocol (https://github.com/ENCODE-DCC/ATAC-seq-pipeline) to obtain the filtered reads and called peaks using MACS2. The filtered bam files from both scATAC-seq and bulk ATAC-seq were converted into normalized bigwig files using deepTools2(*39*). When we aggregated similar neighbor scATAC-seq signals, we adopted the bigWigMerge tool from UCSC website and then converted the bedGraph file into bigwig file using customed script.

Bulk ATAC-seq and 35bp uniqueness mapability signal values were also binned to 1000bp with loci consistent with each ChIP-seq region.

### Data preparation for machine learning

#### Bulk data

The bulk data is prepared as follows. Each bin is one-hot encoded in a 4 x 1000 feature vector *S*. The feature vector *S* is concatenated with the 2 x 1000 ATAC-seq feature vector *A* and the 2 x 1000 mapability feature vector *U*, which refers to the uniqueness of a 35bp subsequence on the positive strand starting at a particular base, to form the input feature vector *S_Bulk_*.

#### Single cell data

The single cell input feature vector *S_SC_* is prepared similarly. The feature matrix *S* is now composed of a bin called by scATAC-seq data. We define the feature vector *A_sc_*, which is the aggregated(*40*)scATAC-seq input data, *A_sc_* is a feature vector identical to *A* in the pre-trained model. The mapability feature vector *U* remains the same.

#### Aggregate scATAC-seq data

The aggregated scATAC-seq data is computed via calculating cell-cell similarity using scATAC-seq binarized cell-peak count matrix. We adopted cisTopic to calculate low dimensional cell-topic latent feature and used cosine similarity to calculate similarity between it and other cells. For each cell, we considered its most 100 similar neighbor cells and aggregated their signals together as its aggregated scATAC-seq data (Supplementary Fig S7).

### Training and prediction

#### Calibration on the TF binding prediction

To train our pretrained models, we chose datasets from three different cell lines. For the TFs that are only present in the dataset from one cell line, those TF outputs were directly used to represent the final TF prediction. If the same TF appeared in multiple cell lines, we calculated the probability of intersecting peaks between called peaks in the single cell dataset and called peaks of each bulk dataset separately. scFAN predicted these TFs on all the three models but only chose one model result whose corresponding cell line has the highest matched probability to the single cell and used its result to represent TF binding.

#### Deep learning calculations

We input the input feature vectors, *S_Bulk_* for the pre-trained model or *S_SC_* for single cell prediction, to a three layer 2D convolutional neural network to extract the feature map. Two fully connected layers are connected to the output feature map, the output of which is passed to a sigmoid function to obtain the prediction of TF binding. Three different pre-trained models were trained on bulk data *S_Bulk_* from three different cell lines (GM12878, K562 and H1ESC). Each model was optimized using the Adam algorithm(*41*), and then individually used to predict TF binding on the single cell data *S_SC_*. The overall deep learning framework is shown in Fig S1. After that, all the TF binding predictions were merged using a calibrated method. For TF exists in multiple cell lines, the most similar cell line was selected and its TF prediction result was chosen, for TF appears only in one unique cell line, its result was used as final result (see details in Supplementary Fig S8).

Our convolution calculation can be defined as follows:

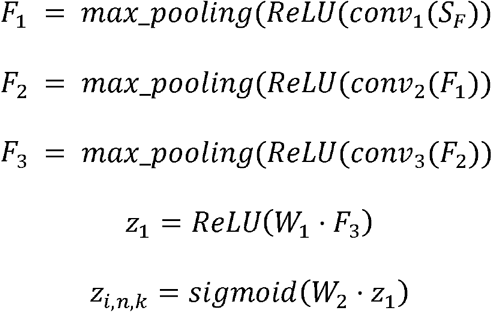

where *S_F_* is either *S_Bulk_* or *S_SC_* and *F*_1_, *F*_2_, *F*_3_, *z*_1_ denote the feature maps of each convolutional layer and the output of the first fully-connected layer. *W*_1_, *W*_2_ refer to weight matrix of the two fully-connected layers. The final output of the network is the probability *z* of TF *k* binding to a peak *n* in each cell *i*, i.e. *z_i,n,k_*, where *k* ∈ 1 ⋯ *M*, *M* is total number of TFs of each cell from the pre-trained model and *n* ∈ 1 ⋯ *N* with *N* being the total number of peaks per cell.

#### Partition choice

Our pre-trained model was restricted to the same dataset partition choice as in Quang *et al*(*8*) for H1-ESC and K562: chromosomes 1, 8, and 21 were used for testing, chromosome 11 was used for evaluation and the remaining chromosomes were used for training (chromosome Y was excluded). The DREAM Challenge dataset for GM12878 cells didn’t include chromosomes 1, 8 and 21, so chromosome 11 was used for both evaluation and testing.

#### TF activity score

Here we select the top two potential predicted TF motifs of each peak and aggregate all the predicted TFs of all the peaks in each cell, then we normalize the value by calculating the probability across all peaks within a cell, which can be defined as the activity score *pc_i,k_* for TF *k* in cell *i*, it was shown as follows:

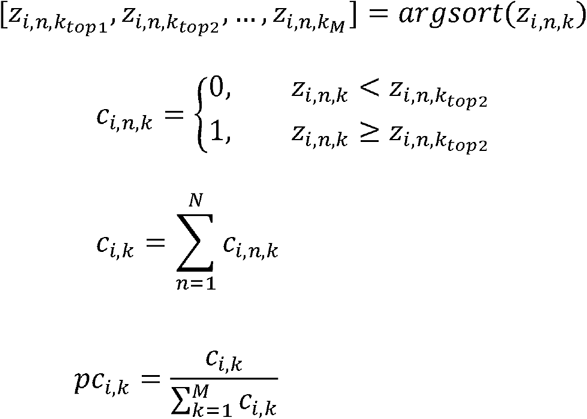

### Cell clustering

The activity score is then used to cluster cells. For clustering convenience, we simply concatenated all the TFs activity score results by column from all three models for all the cells, we did not cut those columns that had repetitive TFs shared by the three models to reduce processing time because dimensionality reduction could perform the same procedure. KNN clustering was then performed and t-SNE plot was drawn using the concatenated feature. To filter the low-quality cells, we set the threshold of the fraction of total read counts per total number of peaks per cell to be 0.05 and also set the threshold of total read counts of each cell to be at least 1000 (Supplementary Fig S2). The scFAN pipeline is shown in Figure 1a.

### Methods comparison

There are several recently published models, scABC(*25*), cisTopic(*26*), Cicero(*28*), SCALE(*27*), Brockman(*29*) and ChromVAR(*14*) that are also designed to cluster single cells based on scATAC-seq data. The first four methods all work with peak-by-cell binarized read count matrix. For example, scABC uses the read count matrix to cluster cells via a weighted K-medoids clustering algorithm. cisTopic adopts Latent Dirichlet Allocation (LDA) to convert the read count matrix into a topic-cell low dimensional matrix, which is further used to clustering cells. Cicero applies Latent Semantic Indexing (LSI) to reduce the high-dimensional matrix into low-dimensional matrix similar to cisTopic. SCALE is a VAE based deep learning model that utilizes gaussian mixture model to initialize and model the cell clusters using binarized peak-cell matrix and uses the latent features to cluster the cells. ChromVAR is based on scATAC-seq read counts and motifs in every peak: single cell-read count matrix and corrected peak-motif matched binary matrix are combined to calculate bias-corrected deviation and z-score matrix. The “corrected” z-score matrix is used to cluster each individual cell. Brockman uses adopted peaks to calculate k-mer frequency within each sample cell, generating over 1000 kinds of k-mer frequency vectors of each cell and uses the combined matrix to cluster the cells. We also adopted raw binarized matrix to directly cluster the cells as benchmark. We used Adjusted Rand Index (ARI), Normalized Mutual Information (NMI) and V-measure score to quantitatively measure the clustering performance of these methods. We determined every cell labels from each method using Euclidean distance and Aggregative clustering based on t-SNE projections of each method, which are the low dimensional t-SNE embedding matrices from scABC, cisTopics, Cicero, and SCALE, k-mer t-SNE embedding matrix from Brockman, the motif correlation t-SNE embedding matrix from chromVAR, and the TF appearance probability t-SNE embedding matrix from our model.

## Availability of Data and Materials

### Data

The bulk ATAC-seq GM12878, H1-ESC and K562 datasets are available from GSE47753, GSE70482 and GSE85330, respectively. The scATAC-seq datasets are available from GSE65360 and GSE74310. The ChIP-seq GM12878 dataset and the K562 and H1-ESC dataset are available from the ENCODE-DREAM Challenge dataset and Li et al(*42*) respectively. The pre-processed datasets are available upon request.

### Software

scFAN is implemented in Python 2.7 based on the Keras library and the model was trained on a NVIDIA TiTan Xp GPU. The code and pre-trained models are freely available at https://github.com/sperfu/scFAN.

## Acknowledgement

We would also like to acknowledge support from NVIDIA.

## Funding

This work was supported by National Natural Science Foundation of China (Grant Numbers 61872288) and China Scholarship Council (to L.F.), NSF grants DMS1763272 and IIS1715017, a grant from the Simons Foundation (594598, QN), and NIH grants U01AR073159 and R01GM123731.

## Author contributions

X.X. conceived the project. L.F. and L.Z. conducted the research. Q.N., Q.P and X.X. supervised the research. L.F., E.D., Q.N., and X.X. contributed to the writing of the manuscript.

## Ethics declarations

### Competing interests

The authors declare no competing interests.

