## Supplemental file 1 for "Predicting transcription factor binding in single cells through deep learning"

### Supplementary file 1

#### Predicting transcription factor binding in single cells through deep learning

Laiyi Fu[1,2], Lihua Zhang[3,5], Emmanuel Dollinger[3,4,5,6], Qinke Peng[1], Qing Nie[3,4,5,6*],

Xiaohui Xie[2,5,6*]

1 Systems Engineering Institute, School of Electronic and Information Engineering, Xi'an Jiaotong University, Xi'an, Shannxi, 710049, China 2 Department of Computer Science, 3 Department of Mathematics, 4 Department of Developmental and Cell Biology, 5 NSF-Simons Center for Multiscale Cell Fate Research, 6 Center for Complex Biological Systems, University of California, Irvine, CA, 92697, USA

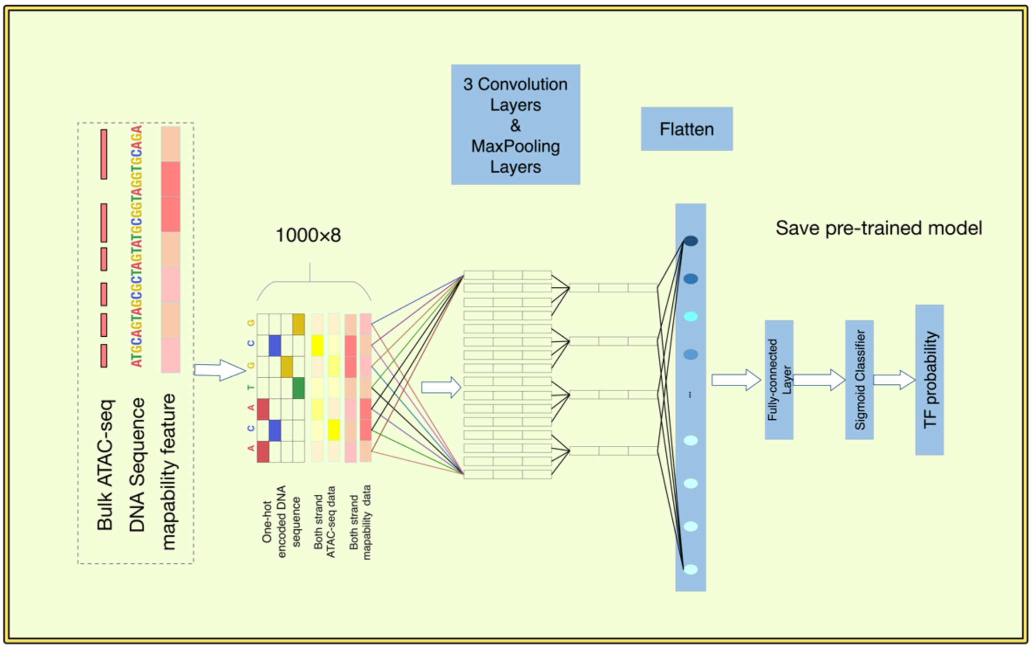


Figure S1: **Deep learning model in scFAN scheme.** The figure shows the overall scheme of the pre-trained model in scFAN. We adopt 3 convolution layers with corresponding Max-pooling layers to extract features, then concatenate the feature map with two fully-connected layers and finally get the output TF probability using Sigmoid function.

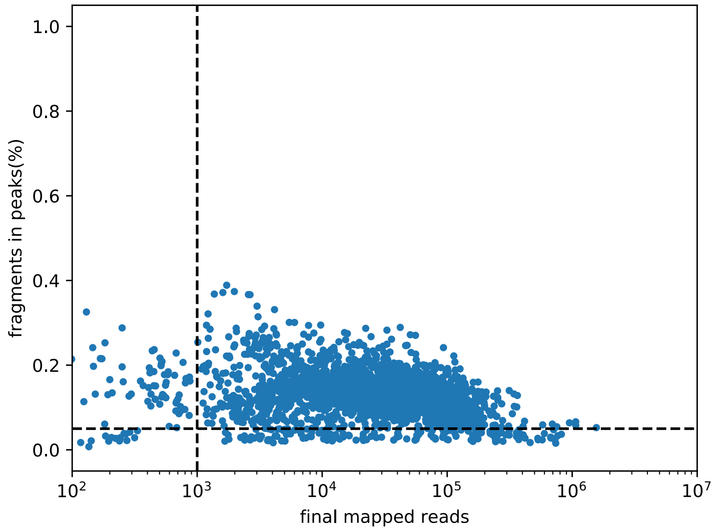


Figure S2: **Single Cell filtering according to read and fragment counts.** The dash lines show the threshold of filtering cells. We set the threshold of the fraction of total read counts per total number of peaks per cell to be 0.05 and also set the threshold of total read counts of each cell to be at least 1000.

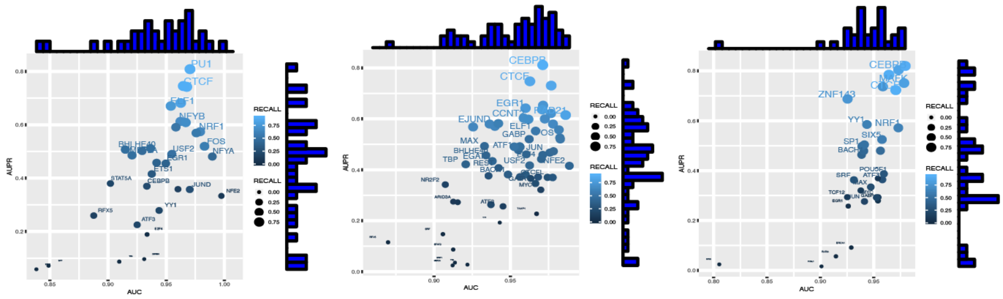


(a) GM12878 based (b) K562 based (c) H1ESC based

Figure S3: **TF binding prediction performance in the pre-trained model using bulk data.** The figure shows the TF prediction performance (AUC, AUPR, and RECALL values) of scFAN implemented by three cell lines based bulk data. X-axis is the AUC value, Y-axis is the AUPR value and the different RECALL value is shown in the lightness of color and the size of the dot. We also draw the histogram plot of the AUC and AUPR along the x-axis and y-axis.

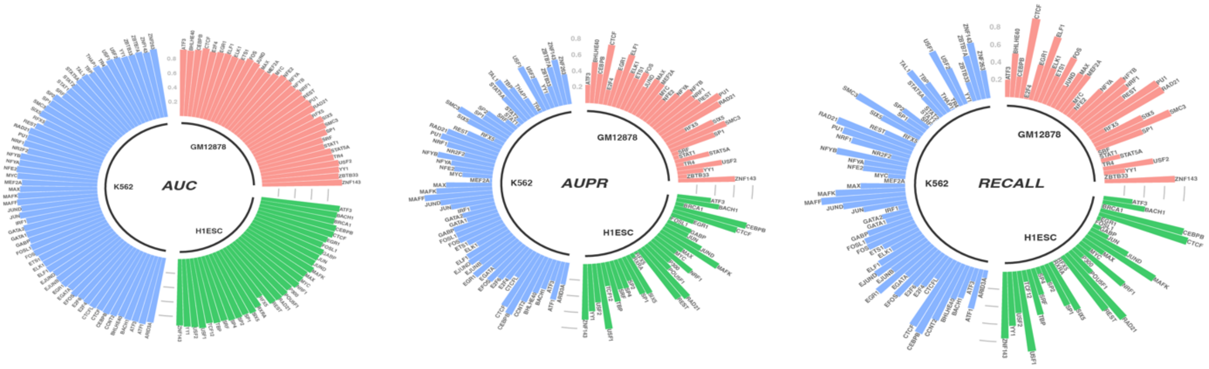


Figure S4: **Each TF prediction performance of in all the three cell lines.** The figure shows each TF prediction performance(AUC, AUPR, and RECALL values) of three cell lines (GM12878/K562/H1ESC) using scFAN pre-trained model. The higher the bar is, the larger the value is

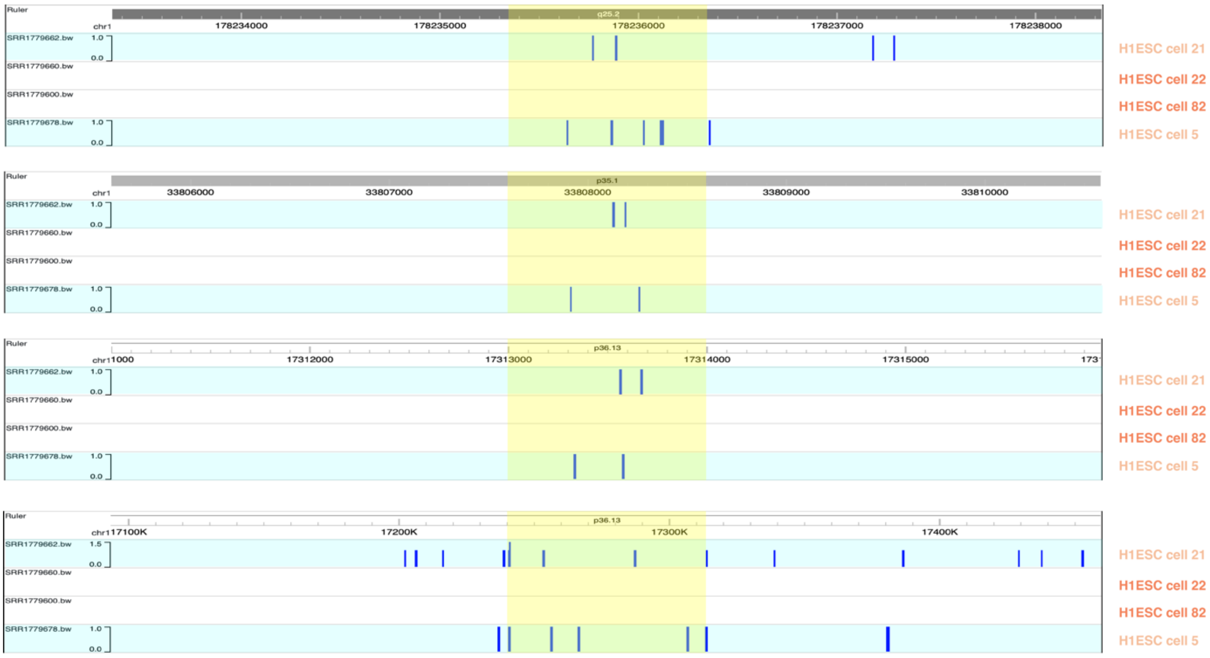


Figure S5: **The raw scATAC-seq signal across four H1ESC cells in chromosome 1.** The figure shows the appearance raw signal of scATAC-seq data in the predicted regions (from four different regions in chromosome 1) that are bind with CTCF across four H1ESC cells. Each blue panel (the first row and the last row) indicate one H1ESC cell that shows more active score (more scATAC-seq signal) than the white panels (the second row and the third row), which results in the separation on the clustering of H1ESC cells.

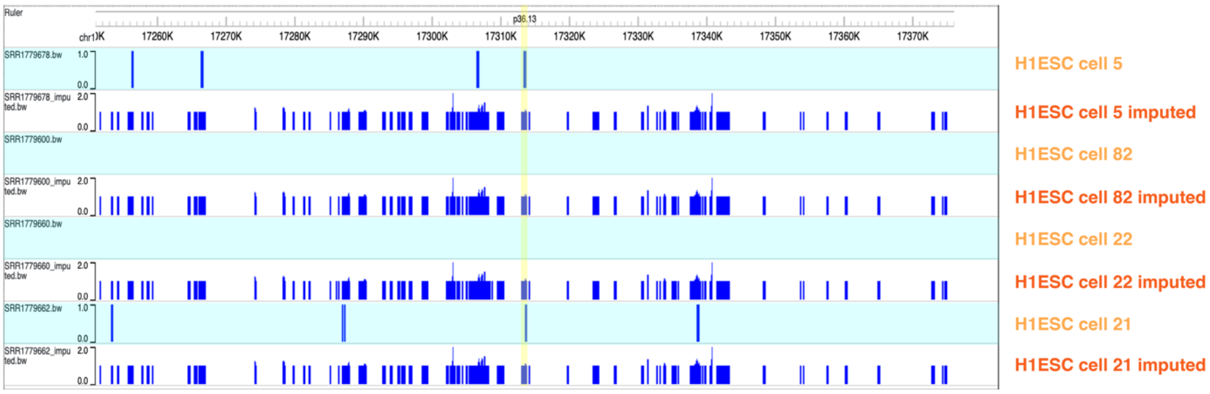


Figure S6: **The imputed ATAC-seq and scATAC-seq signal from the same regions.** Four cells are chosen. The data in blue panel is raw scATAC-seq data and the white panel is the imputed scATAC-seq data. After the imputation, those regions that are originally missing some signal were recovered, which lets the model able to predict TF in these regions more accurately and further clusters these cells together.

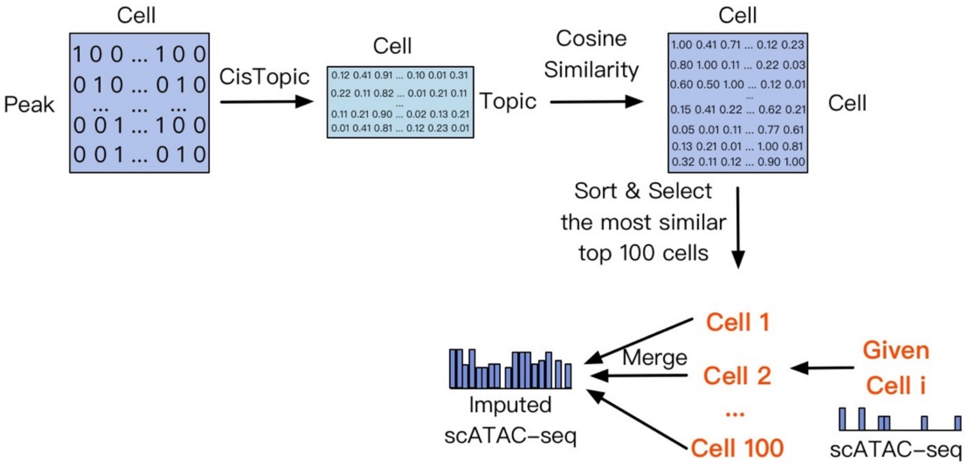


Figure S7: **The imputation procedure of scATAC-seq data.** The figure shows the procedure of implementing scATAC-seq imputation using binarized peak-cell count matrix. We used cisTopic to reduce the original high-dimensional matrix into low-dimensional latent space(topic-cell) matrix, which can be further used to calculate similarity matrix between each cell. Given a single cell i, we sorted the similarity matrix and select the most similar 100 cells to cell i, and merge their signal into one file using bigWigMerge tool to get the final imputed scATAC-seq data with respect to cell i.

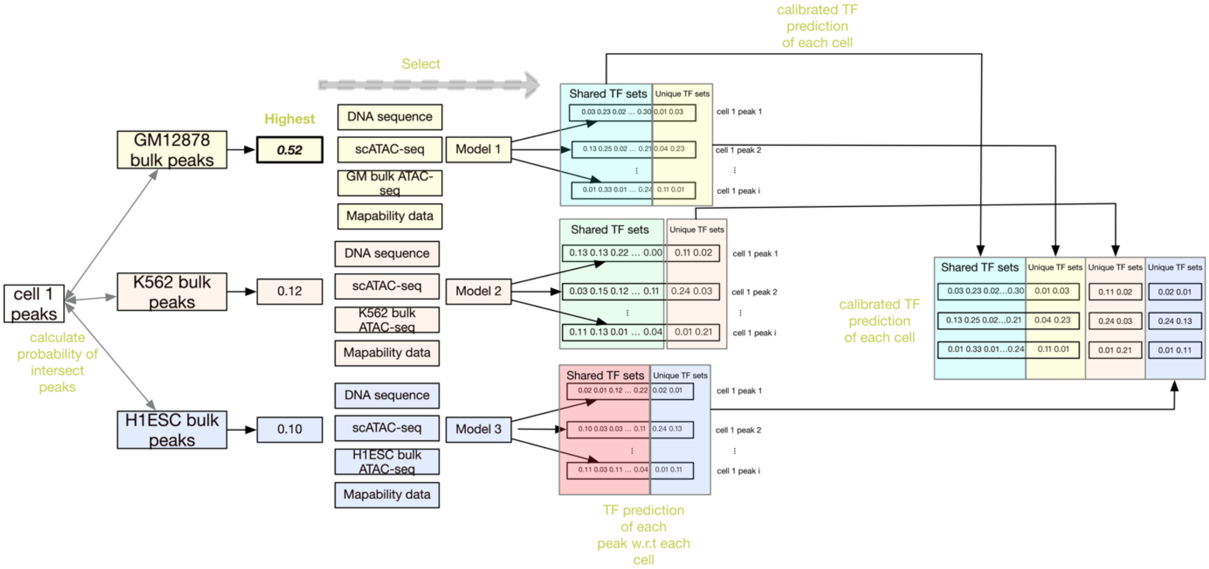


Figure S8: **The calibration of TF binding prediction in one single cell.** The figure shows the calibration of TF binding prediction from three different models. All the peaks in one single cell were intersected with bulk peaks from three cell line. The probability of the intersecting number in single cell was calculated and the highest one was selected as the most likely matched cell line and chosen as the representation of all the shared TFs by three models. All the unique TFs in three models were kept separately. After that, the calibrated TF prediction of each cell was obtained.

Table S1: **Partial summay of scFAN cross-cell performance on bulk data.** From the table we could see that our model could capture different TFs from different cell lines.

| TF name | Cell line | AUC | AUPR | RECALL |
|---|---|---|---|---|
| CTCF | GM12878 | 0.968 | 0.742 | 0.919 |
| ELK1 | GM12878 | 0.954 | 0.670 | 0.765 |
| PU1 | GM12878 | 0.970 | 0.808 | 0.911 |
| RAD21 | GM12878 | 0.965 | 0.745 | 0.912 |
| SMC3 | GM12878 | 0.963 | 0.682 | 0.845 |
| CTCF | K562 | 0.963 | 0.748 | 0.855 |
| CEBPB | K562 | 0.971 | 0.811 | 0.904 |
| MAFF | K562 | 0.977 | 0.731 | 0.848 |
| RAD21 | K562 | 0.986 | 0.615 | 0.829 |
| EGR1 | K562 | 0.960 | 0.642 | 0.762 |
| JUND | H1ESC | 0.941 | 0.479 | 0.463 |
| CEBPB | H1ESC | 0.973 | 0.804 | 0.865 |
| CTCF | H1ESC | 0.971 | 0.723 | 0.937 |
| MAFK | H1ESC | 0.978 | 0.728 | 0.840 |
| USF1 | H1ESC | 0.979 | 0.820 | 0.925 |
